# The complex relationship between effort and heart rate: a hint from dynamical analysis

**DOI:** 10.1101/2020.03.01.971929

**Authors:** D. Mongin, C. Chabert, A. Uribe-Caparros, J. F. Vico Guzmán, O. Hue, J. R. Alvero-Cruz, D. S. Courvoisier

## Abstract

Heart rate during effort test has been previously successfully adjusted with a simple first order differential equation with constant coefficients driven by the body power expenditure. Although producing proper estimation and yielding pertinent indices to analyze such measurement, this approach suffers from its inability to model the saturation of the heart rate increase at high power expenditure and the change of heart rate equilibrium after effort. The objective of the present study is to improve this model by considering that the amplitude of the heart rate response to effort (gain) depends on the power expenditure value. Therefore, heart rate and oxygen consumption of 30 amateur athletes were measured while they performed a maximum graded treadmill effort test. The proposed model was able to predict 99% of the measured heart rate variance during exercise. The gains estimated at the different power expenditures were constant but noisy before the first ventilatory threshold, stable and decreasing slightly with power increase between the two ventilatory thresholds, before decreasing in a more pronounced manner after the second ventilatory threshold. The slope of the decrease of heart rate gain with power expenditure was correlated with the deflection angle of the heart rate performance curve and with the maximum oxygen consumption. These results reflect the changes of metabolic energy systems at play during the effort test and are consistent with the analysis of the heart rate performance curve given by the Conconi method, thus validating our new approach to analyze heart rate during effort test.

## Introduction

Measurement of heart rate (HR) during effort test is a key measurement to assess physical performance ^1^ and is everyday more accessible, thanks to the development of mobile measurement device ^2^. In addition to the calculation of the heart resting rate (HRR) and the HR reserve ^3,4^, the HR measurement curve obtained during a maximal graded exercise test is also been employed to determine the lactic accumulation threshold ^5^. This method, known as the Conconi method, link the lactic accumulation threshold with the HR dynamic change at high energy expenditure, appearing as an inflection point in the HR performance curve. Attractive by its simplicity, this approach suffers from several drawbacks, including a variable reliability ^6,7^, the existence of HR performance curve having no inflection points ^8–10^, and a lack of accuracy ^11,12^.

Recent developments of dynamical analysis based on first order differential equation have proven to very accurately reproduce the dynamics of oxygen consumption (VO_2_) and HR during any effort type ^13,14^, and to produce indices sensible to physical performance and physical fitness ^14^. Among these indices, the HR gain, that is the proportionality between the effort and the HR increase, was linked with most of the performance indicators. Despite these promising results, the dynamical approach currently used suffers from the fact that it supposes a constant gain and a constant equilibrium value over the whole effort test, being thus unable to model HR dynamic changes at high energy expenditure ^15^. Indeed, at high level of effort, the heart rate cannot increase as much as it did for low level of effort^15^. In addition, HR is known to first reach a HR value higher than its resting value just after the effort, as it decreases back to its resting value on a longer time scale due to the reduction of blood volume (i.e. dehydration), the evacuation of the heat accumulated during the muscular contractions, or the over-activation of the sympathetic system during exercise ^16–18^. A dynamical model allowing the gain to vary with the power spent during the exercise test would allow to model more properly the HR dynamics and its allostasis (i.e., return to a higher resting heart rate than before the test) during effort and would thus allow to quantify the HR dynamics change occurring during the test.

We propose in this article to apply to HR measurements an extension of the dynamical model proposed previously to account for gain change with energy expenditure along grading exercise tests. The quality of the estimation provided by this model will be compared to the previous one. The quantification of the HR dynamic change stemming from this analysis will be compared with the Conconi method and linked with standard performance indices.

## Methods

### Population

The population studied consisted in 30 amateur athletes (28 males, 2 females, Age = 36 ± 8 years). Table 1 summarize the physiological characteristics of this population. The mean fat percentage is low, and the maximum aerobic O_2_ consumption is high with low standard deviation, indicating an ensemble of well-trained persons.

**Table 1:** physiological characteristics and standard performance indices of the population

|  |  |
| --- | --- |
| **Number of individuals** | **30** |
| **Age (years)** | 36.07 (8.19) |
| **Height (cm)** | 174.26 (6.82) |
| **Weight (kg)** | 72.52 (9.70) |
| **BMI** | 23.83 (2.39) |
| **Sex (N male)** | 28 (93.3%) |
| **Fat (%)** | 16.15 (2.65) |
| **VO<sub>2</sub> max (mL/kg/min)** | 50.18 (6.16) |
| **Maximal Speed (km/h)** | 17.71 (2.07) |
| **HR max (beat/min)** | 182.33 (7.83) |
| **VT1 (km/h)** | 9.23 (1.52) |
| **VT2 (km/h)** | 13.20 (1.88) |
| **HRR (beat/min)** | 36.07 (8.91) |

Note: VO<sub>2</sub> max: maximal aerobic capacities, VT1 and VT2: speed at the first ventilator and second ventilatory transition, HR max: maximum HR; HRR: Heart Resting Rate.

### Maximal effort test

The athletes performed graded exercise on a PowerJog J series treadmill connected to a CPX MedGraphics gas analyzer system (Medical Graphics, St Paul, MN, USA) with measurement of respiratory parameters of which the total volume exhausted (VE), oxygen consumption (VO_2_), carbon dioxide consumption (VCO_2_), and HR with a 12 lead ECG. Measurement are taken cycle to cycle. The stress test consisted of an 8-10 min warm up period of 5 km.h^−1^ followed by step 1km.h^−1^ by minute speed increase until maximum effort was reached. Power developed during the effort test was calculated using formula described by the American College of Sport Medicine (ACSM) to determine an approximate VO_2_ of runners ^19^ associated to the Hawley and Noakes equation linking oxygen consumption to mechanical power ^20^.

### Standard performance indices

The HRR calculated is the standard HRR60, which is the difference between the HR at the onset of the recovery and the HR 60 seconds later. The ventilatory thresholds 1 (VT1) and 2 (VT2) are calculated using the Wasserman method using the minute ventilation (VE)/VO_2_ for determining VT1 and VE/VCO_2_ for VT2 ^21^.

Concerning the Conconi method, we focused on the deflection degree of the HR curve as presented in Hoffman’s and coauthors work ^9^. Shortly, calculating this quantification of the HR curve deflection consists in fitting the HR curve with a second-degree polynomial of time for powers between the power at first ventilatory transition (PVT1) and the maximal power, in order to calculate the tangent slopes of the HR curve at these two power values (the two tangent k1 and k2 respectively at power equal to PVT1 and P max). This calculation permits to calculate the deflection degree of the HR curve:

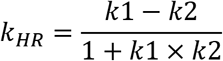

Following ^9^, *k*_*HR*_ > 0.1 is considered to be a normal deflection, kHR < 0 is an inverse deflection and 0 < *k*_*HR*_ < 0.1 marks an absence of usable deflection point.

### Dynamical analysis

The basis of the dynamical analysis proposed rely on the estimation of the constant coefficients of a first order differential equation (for more details, see ^14^). Shortly, describing the HR dynamics by a first differential equation consists in linking the change in time of HR with its value and the power spend during effort:

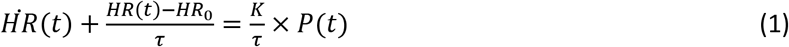

Where 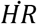 is the time derivative of *HR* (i.e. its instantaneous change over time) and *P*(*t*) the power spent during the effort. Equation 1 describes the dynamics of a self-regulated system with an equilibrium value *HR*_0_, a damping time *τ* and a gain *K* (see Figure 1 left panel). The equilibrium value corresponds in the case of HR to the heart rate resting value *HR*_0_ (in beat/min), and is the HR in the absence of effort. HR described by equation 1 will respond to a constant energy expenditure *P* by increasing from *HR*_0_ to *HR*_0_ + *KP* in an exponential manner, with the damping time *τ* associated being the time needed for HR to reach 63% of its total change. The gain (in beat/min/W) gives the proportionality between an effort increase Δ*P* (in W) and the associated total HR increase Δ*HR* (see Figure 1 left panel):

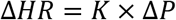

**Figure 1:**
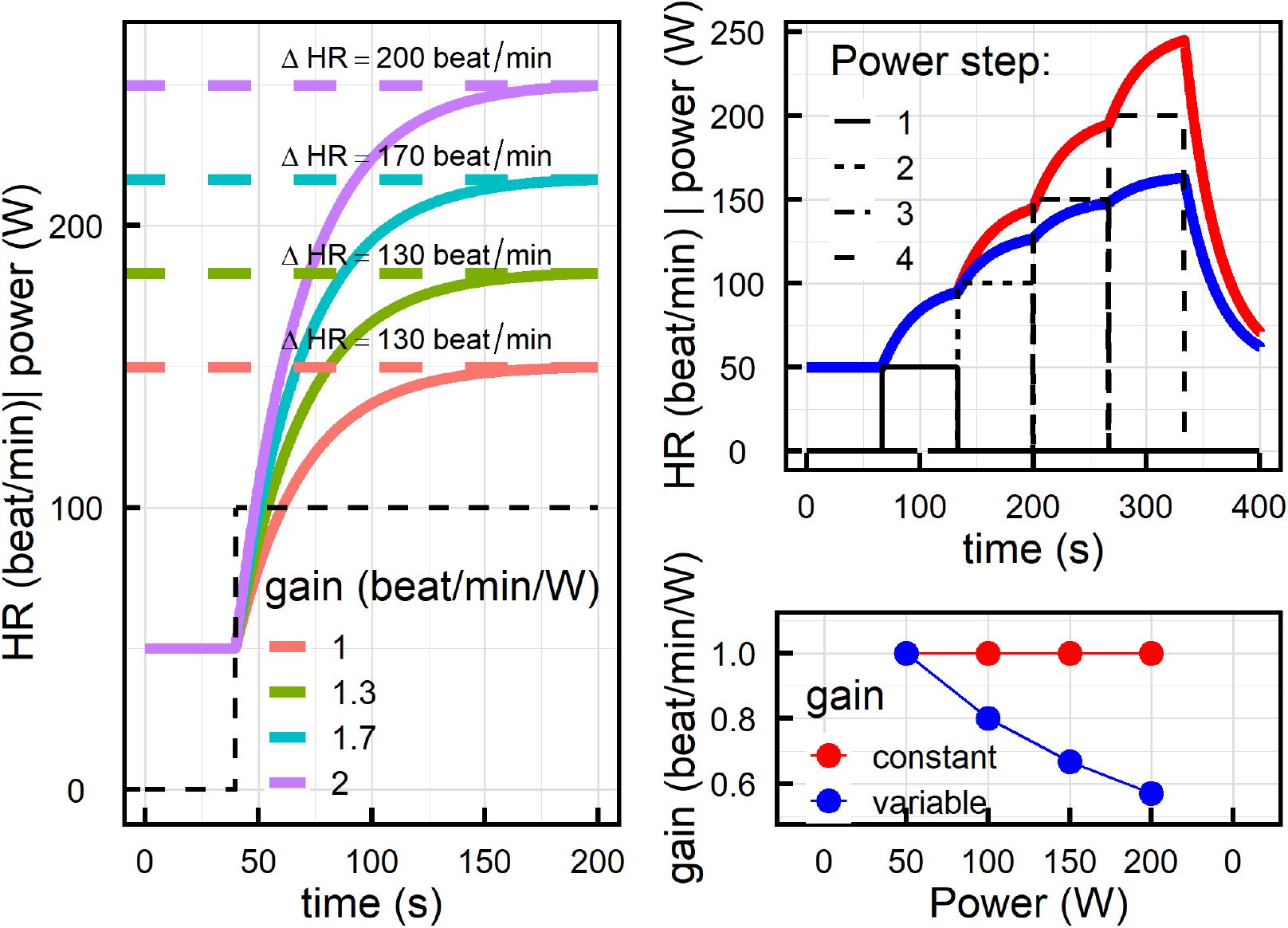
left: HR response following a first differential equation driven by a constant step effort of 100W. Parameter are *τ* = 30*s*, *HR*_0_ = 50 beats/min and four values of gain. Right: HR response following a first differential equation driven by an incremental effort, with *τ* = 30*s*, *HR*_0_ = 50 beats/min with a constant gain of 1 beat/min/W (red curve) and a gain decreasing with power (blue curve).

Determination of these 3 coefficients (resting value, damping time and gain) is performed by first evaluating the HR derivative over multiple points, and then performing a multilevel linear regression between 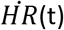, *HR*(*t*) and *P*(*t*).

To account for a power dependent gain, it is possible to decompose the power as a sum of successive power steps, each of them having an associated gain:

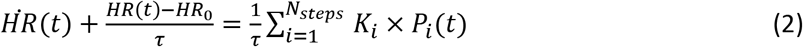

Where the index *i* ranges from 1 to *N*_*steps*_ the total number of power steps considered. The gain *k*_*i*_ is representative of the HR response for the power encompassed by *P*_*i*_(*t*) (see Figure 1 right bottom panel). An example of the HR response to an incremental effort with a decreasing gain compared to the response with a constant gain is presented in the right panel of Figure 1. In the case of a graded exercise, the natural decomposition of the power spend during the test is a sum of constant power steps. In the example depicted in the right panel of Figure 1, the 4 power steps introduced in equation 2 allow to obtain four values of gain for their respective power ranges (right bottom panel of Figure 1).

This method allows at the same time to model a simple allostasis, that is to account for a change of equilibrium value. Indeed, by artificially setting the power of the recovery period to 1W before the estimation of the first order differential equation, the equilibrium value after the effort will be *HR*_0_ + 1 × *k*_*recovery*_, different from *HR*_0_.

### Sensitivity analysis

To ensure that the variation of the HR gain with power was not dependent of the number of steps *N*_*steps*_ considered for the decomposition of the incremental effort test power, we performed the above described dynamical analysis for different values of *N*_*steps*_.

### Statistical analysis

Associations between continuous variables of interest were assessed using Spearman correlation. Associations between gain and power was analysed using mixed effect linear regression. All analyses were performed using R version 3.4.2, the package doremi for the dynamical analysis, lmer and lme4 for the mixed effect linear regression, and the packages data.table, Hmisc and ggplot2 for the data management and statistical indicators.

## Results

Performance of the new dynamical analysis of HR with a variable gain is illustrated for one effort test in Figure 2 (left panels), and compared for the same effort test with the dynamical analysis with a constant (right panels). The estimation of the HR dynamics with a constant gain yields a median R^2^ of 0.91 (interquartile range IQR = [0.89-0.95]), while the new dynamical analysis considering the power as a sum of constant steps yields a median R^2^ of 0.994 (IQR: [0.987- 0.996]) (bottom panels).

**Figure 2:**
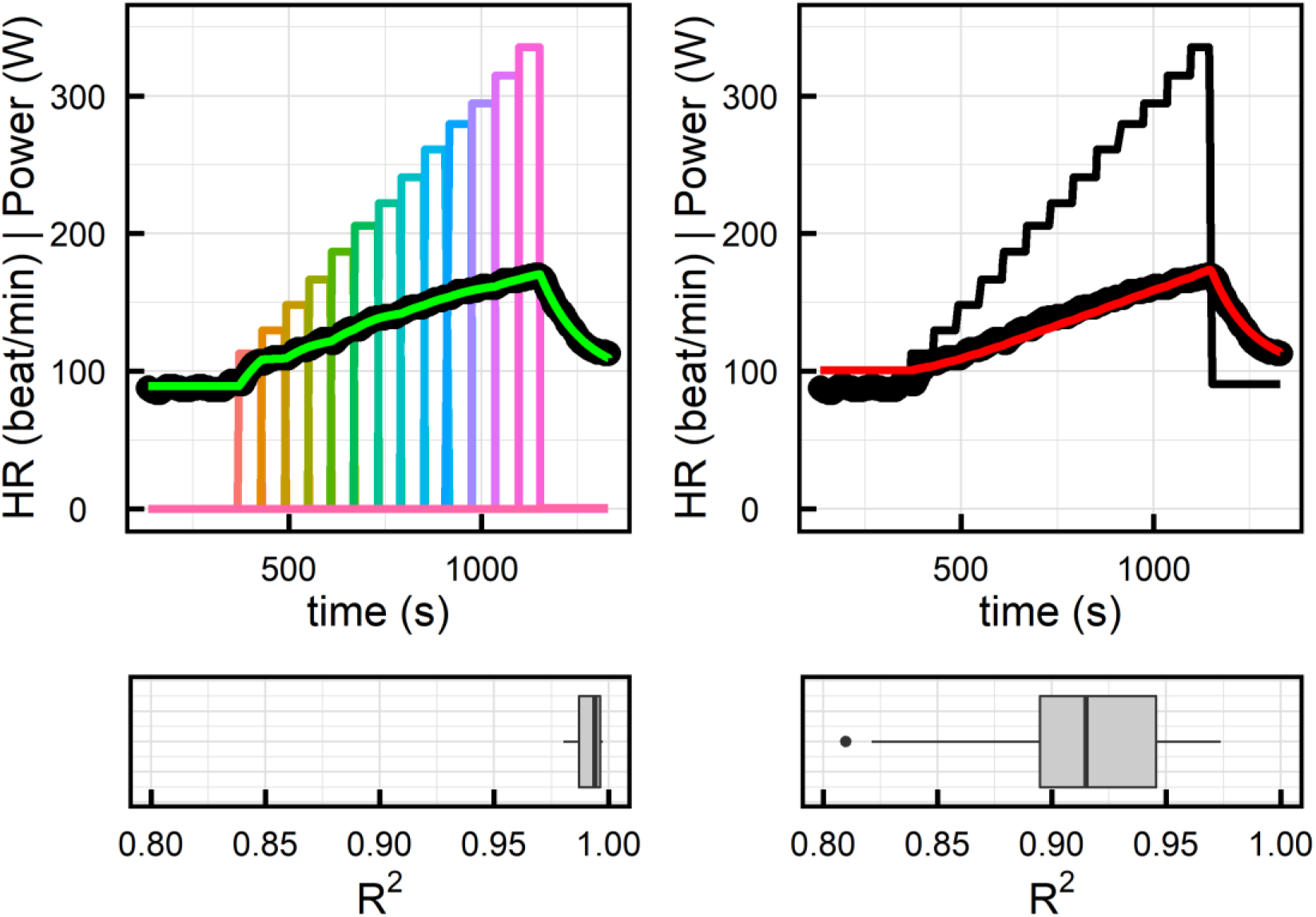
Example of fitted HR curve with a dynamical analysis considering the energy expenditure as one and a constant gain (right panel) or considering the energy expenditure as a sum of constant steps with a gain associated to each step (left panel). The boxplots below the graphs represent the ensemble of the R^2^ value of the estimations.

The variable gain approach allows to estimate an unbiased equilibrium value *HR*_0_, whereas as already previously seen^14^ the constant gain approach tends to overestimate it (see Supplementary Figure 1).

Since the dynamical change of the HR curve is linked to the ventilatory thresholds ^15^, HR gains are described for power below the first threshold, between the two ventilatory thresholds and above the power corresponding to the lactate turn point. A representative measurement of the estimated HR gain in function of the power spent during exercise is displayed on Figure 3, together with the two ventilatory transitions estimated from the ventilatory measurements.

**Figure 3:**
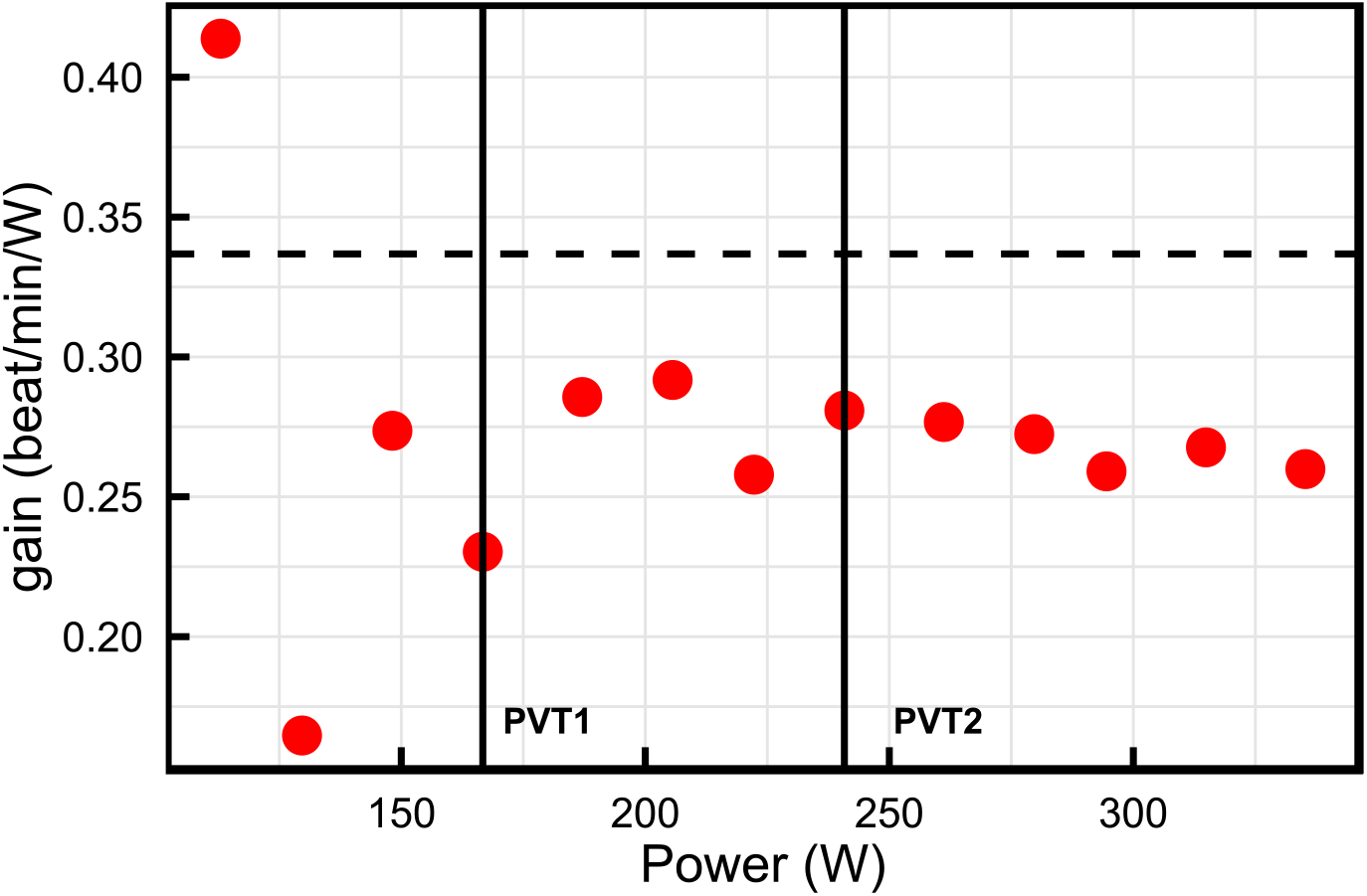
Example of HR gain (dots) for the different power values considered. The two vertical line represent the two ventilatory thresholds, and the horizontal dashed line the value of the constant HR gain.

Using linear mixed models, we examined the trend in gain over time (in this effort test, time and power are interchangeable). Before the first ventilatory transition, the gain varies greatly but with no trend before decreasing slightly between the two transitions and more strongly after the second transition (Table 2).

**Table 2:** Estimated linear trend of HR gain over time, for three power ranges (PVT: power at ventilator transition). N indicates the total number of HR measurements considered for the mixed effect regression. Significance is indicated as follows: *: *p*<0.05; **: *p* < 0.01, ***: *p* < 0.001

| Power range | Slope (beat/min/kW<sup>2</sup>) (sd) | N | R<sup>2</sup> | P value |
| --- | --- | --- | --- | --- |
| Power ≤ PVT1 | -178 (391) | 117 | 0.55 | 0.65 |
| PVT1 ≤ Power < PVT2 | -316 (134) | 130 | 0.80 | 0.02 |
| PVT2 ≤ Power | -528 ( 55) | 146 | 0.94 | <0.001 |

R^2^ illustrates that the gain varies a lot around its mean constant value for low powers, has a reduced variability around its decreasing trend between the two ventilatory thresholds, and stabilize in a more pronounced decrease after the second threshold.

These individual slopes of gain versus power can be used as an indicator of heart performance and function. Correlations and comparisons with other cardiovascular performance indices are given in Table 3.

**Table 3:** Spearman correlation coefficients between various measures of performance

|  | Trend in HR gain |  | HR Constant gain | HRR | HR max |
| --- | --- | --- | --- | --- | --- |
|  | PVT1 < Power < PVT2 | Power > PVT2 |  |  |  |
| HR constant gain | -0.67*** | -0.33 |  |  |  |
| HRR | 0.19 | 0.14 | 0.13 |  |  |
| HR max | -0.38* | -0.01 | 0.15 | -0.16 |  |
| VO2 max | 0.29 | 0.48** | -0.41* | 0.20 | 0.19 |
| Speed max | 0.31 | 0.33 | -0.50** | 0.30 | 0.07 |
| PVT1 | -- | -- | -0.37* | -0.01 | -0.21 |
| PVT2 | -- | -- | -0.70*** | 0.04 | -0.09 |
| k<sub>HR</sub> (Conconi) | -0.28 | -0.57** | 0.32 | 0.07 | 0.21 |

*Note: HRR: Heart Resting Rate, HR max: maximum HR, VO2 max: maximal oxygen consumption, Speed max: maximal speed achieved during the effort test, PVT1 and PVT2: power at the ventilatory transitions; k<sub>HR</sub>: HR curve deflection angle. Significance is indicated as follows: \*: $p < 0.05$ ; \*\*: $p < 0.01$ , \*\*\*: $p < 0.001$*

Between the two ventilatory thresholds, the decrease in gain is strongly associated with higher mean HR gain and moderately associated with higher HR max.

After the second threshold, the decrease rate of HR gain with power is correlated with lower VO_2_ max. In other words, a higher maximal aerobic respiratory capacity is associated with a smaller decrease of the HR gain with power. This correlation is higher and more significant than the one between the constant HR gain and VO_2_ max. The decrease rate of HR gain with power above the lactic accumulation threshold is negatively correlated with the deflection angle of the HR curve measured with the Conconi technique, meaning that stronger deflections of the HR performance curve are associated with stronger decrease of the HR gain with power increase. The slopes of HR gain evolution with power for the two power zones considered are strongly correlated with each other (ρ=0.68, p < 0.001).

### Sensitivity analysis

Decomposition of the effort test power in a fixed number of parts ranging from 4 to 15 did not qualitatively change the results. Specifically, the tendency of the evolution of the gain with power as presented in Table 1 stayed the same, as well as the correlation between the decrease of the HR gain with power and the other indices (Table 3). An example of the estimations of HR obtained by our dynamical analysis for the different *N*_*step*_ considered together with the values of gain estimated are presented in Supplementary Figure S1.

## Discussion

### Main findings

Considering multiple independent power steps in the dynamical analysis of heart rate allowed to study the heart rate dynamical changes occuring during effort tests and to account for the inherent allostasis of the heart functionning during physical exercise. This approach produced an estimation of HR accounting for more than 99% of its observed variation and allowed to estimate the HR response to each independent power steps considered. Before the first ventilatory threshold, the HR gain (i.e. the proportionality between HR increase and effort increase) was independent of energy expenditure and had an important variability. The HR gain started to decrease with power increase between the first and the second ventilatory threholds, and decreased more with power after the second ventilatory threshold. The slope of these decreases was correlated with the constant HR gain between the two ventilatory thresholds, and with VO_2_ max and the HR curve deflection angle for power above the lactic accumulation threshold.

### HR dynamical changes

The HR dynamics captured by the HR gain changes with power can be explained by the various metabolic energy systems at play during the exercise ^22,23^.

At the onset of a physical effort, the energy needed for mechanic workload is first supplied by anaerobic pathways (including phosphagen then glycolysis systems), and progressively replaced by aerobic pathways after a few minutes of exercise ^23,24^. In most of the graded exercise used, the fast intensity increase leads to the significant involvement of the main energetic pathways below the first ventilatory threshold. In this situation, a non negligeable part of the energy supply does not rely on exogenous oxygen nor substrates provided by blood compartment, causing the important variability of the HR gain measured by our analysis before VT1. Futhermore, heart rate adaptation at the onset of exercise is driven by both nervous and endocrine systems that have differents response inertia, participating to the heart rate variability measured in this study at the beginning of exercise ^25,26^.

Between VT1 and VT2, energy expanditure is mainly supplied by oxidation of substrates in mitochondria. In this range of intensity, odxydated subtrates shift from lipids to glucose, the only subrate used at high intensity, and lactate starts to progressively accumulate faster than it can be eliminated ^27^. Despite the better yield of glucose oxydation, a slight decrease of the HR gain with power is observed at this intensity range, which may be due to the negative effects of progressive lactate accumulation in muscle ^28^. Lactate may also explain the correlation observed between the mean HR gain and the HR gain decrease. The fact that athletes with higher mean HR gain, i.e. higher heart rate increase for a given energy increase, have a stronger decrease of the HR gain with power, may be due to a better lactate management by athletes with a lower HR gain. This result is in line with current litterature that describes the relation between performance capacity and development of lactate countermeasures ^29^. Nonetheless, the almost exclusive participation of the aerobic pathway leads to the more stable HR gain observed, despite the physiological disturbances induced by lactate.

After VT2, aerobic then anaerobic glycolysis are the main contributors to the energy demand. At such intensity, all the physiological machinery is working to maintain mechanic workload whithout variations induced by metabolic energy systems inertia or lipid to glucid oxydation shift, causing the HR gain to become even more stable. However, the lactate countermeasures are overloaded, leading to a fast increase of lactate concentration and a decrease of the muscle pH that sharply affects muscle performance^30^. The HR gain thus starts to decrease in a more prononced manner with power increase, expressing the physiological limit to effort increase. This change observed in HR dynamics is related with aerobic energy supply capacity, as seen by the negative correlation between the decrease of HR gain and VO_2_max. Athletes with higher VO_2_max have a lower decrease of HR gain with power, i.e. a better ability to counteract blood lactate accumulation and maintain a better cardiorespiratory response to the increasing energy demand ^29^.

### Comparison with Conconi

The main objective of Conconi test is to determine VT2. Our analysis focuses more in the quantification of the HR dyncamical change. Our analysis provides a measurement similar to the deflection degree measured directly on the HR curve, as can be seen with the correlation between the HR gain decrease after the second ventilatory threshold and the deflection degree of the HR performance curve. But contrary to this measurement, ^15,31^ our approach is not dependent on the protocol ^13,14^. The fact that the HR gain starts decreasing between the first ventilatory and the second ventilatory threshold could explain the difficulties encounter to systematically observe the deflection of the HR performance curve with the Conconi approach.

### Comparison with dynamical approach with constant gain

While the dynamical analysis considering a constant gain gives an average measurement of the HR response to effort and is representative of the overall athelte performance, the dynamical analysis proposed in this study provides better HR estimate than the previous one, mainly due to its ability to model allostasis. It allows a deeper insight into HR dynamics with a non invasive way to quantify the HR dynamic change during lactate production and accumulation, which constitutes a new and pertinent heart characterisation.

### Strengths and limitations

Strengh of the study is the high estimation performance that the model is capable of, as well as the consistence with known physiological behaviors. The dynamical analysis is applicable to any kind of protocol. In addition, well-known indices were also calculated, to allow head-to-head comparison with the new method used.

There are two main limitations of this study. First, the sample size is relatively small and the sample is only composed of amateur athletes. Thus, the sample is not very heterogeneous. Thus, further studies should consider various effort test, and diverse populations.. The second limitation is the lack of lactate measurements, which would have allowed a direct testing of the association between HR gain and lactate management.

## Conclusion

Dynamical analysis of heart rate with multiple independent power steps is a promising method to analyse effort tests. It provides a highly precise modeling of heart rate dynamics (R^2^ > 0.99) with only a few parameters. HR gain and change in HR gain in particular yield good insights into the physiological underpinings of athletes performance during effort test.

## ACKNOWLEDGEMENTS

D.M. work was supported by the SNSF scientific exchange grant IZSEZ0_183540 and the ICARUS SNSF fund 100019_166010.

A.U.C work was supported by the ICARUS SNSF fund 100019_166010.

C.C. and O.H. were supported by the « Programme Opérationnel - Fonds Européen de Développement Régional »(PO-FEDER) under grant 2015-FED-213, GP0008037.

---

**Supplementary Figure S1:**
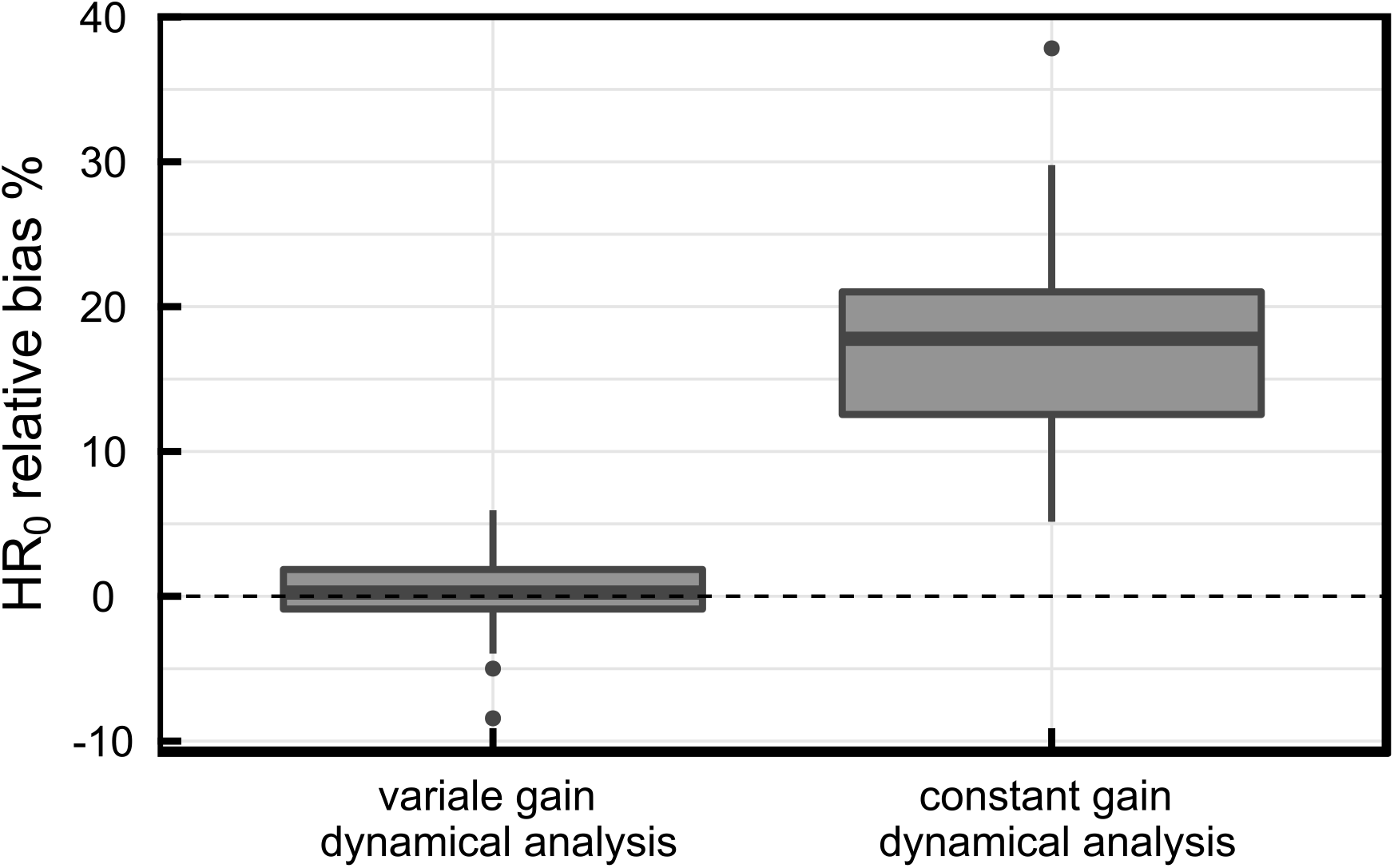
Relative bias of the estimated equilibrium value compared to the actual mean HR value before the beginning of the incremental effort test, for the two analysis.

**Supplementary Figure S2:**
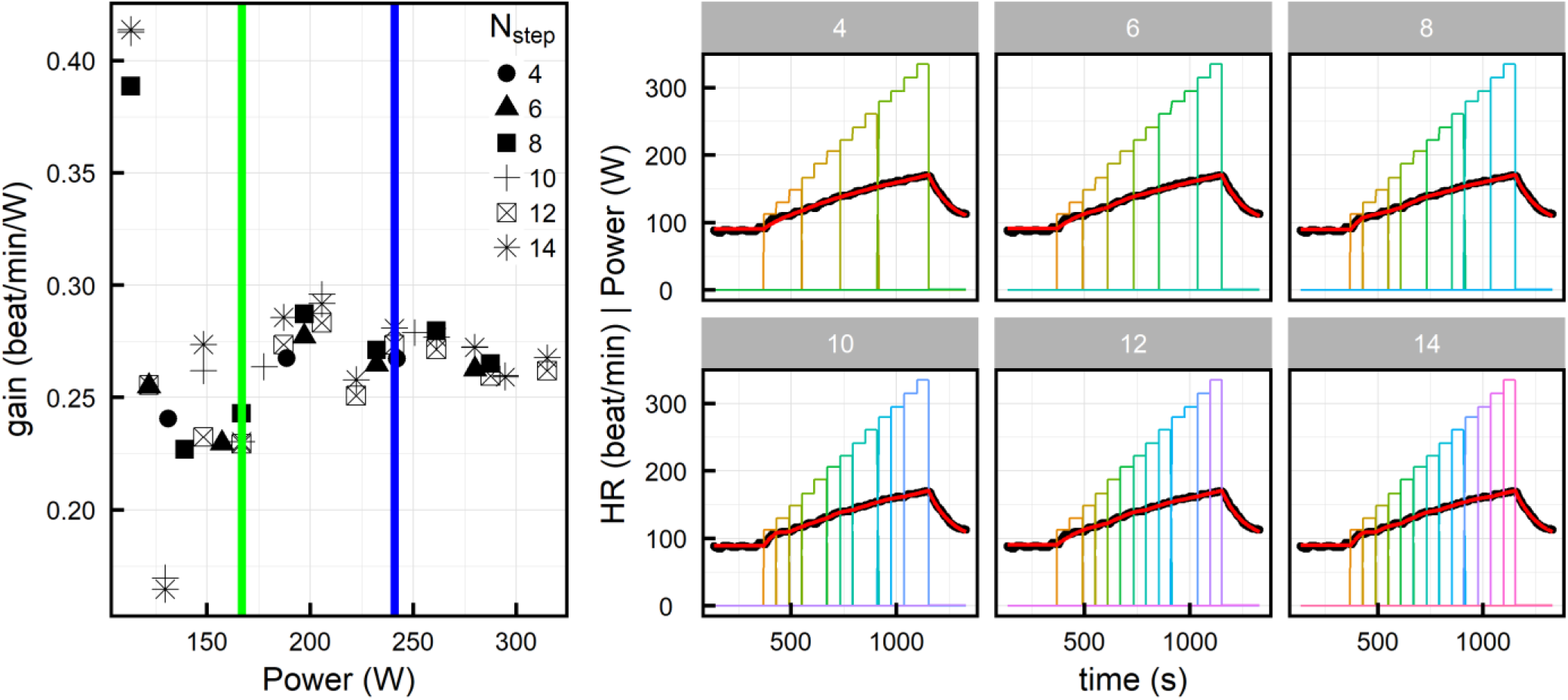
right: HR estimation (red line) compared to the experimental HR (black dots) produced by our dynamical analysis for 6 different energy expenditure decomposition (coloured solid line, for *N_step_* = *4, 6, 8, 10, 12, 14*). Left: HR gain estimated from the different analysis presented in the right panel.

## References

1. Bellenger CR, Fuller JT, Thomson RL, Davison K, Robertson EY, Buckley JD. Monitoring Athletic Training Status Through Autonomic Heart Rate Regulation: A Systematic Review and Meta-Analysis. Sports Med. 2016;46(10):1461–1486. doi:10.1007/s40279-016-0484-2

2. Clark CCT, Barnes CM, Stratton G, McNarry MA, Mackintosh KA, Summers HD. A Review of Emerging Analytical Techniques for Objective Physical Activity Measurement in Humans. Sports Med. 2017;47(3):439–447. doi:10.1007/s40279-016-0585-y

3. Bunc V, Heller J, Leso J. Kinetics of heart rate responses to exercise. Journal of Sports Sciences. 1988;6(1):39–48. doi:10.1080/02640418808729792

4. Ludwig M, Hoffmann K, Endler S, Asteroth A, Wiemeyer J. Measurement, Prediction, and Control of Individual Heart Rate Responses to Exercise-Basics and Options for Wearable Devices. Front Physiol. 2018;9:778. doi:10.3389/fphys.2018.00778

5. Conconi F, Ferrari M, Ziglio PG, Droghetti P, Codeca L. Determination of the anaerobic threshold by a noninvasive field test in runners. J Appl Physiol Respir Environ Exerc Physiol. 1982;52(4):869–873. doi:10.1152/jappl.1982.52.4.869

6. Cabo JV, Martinez-Camblor P, del Valle M. Validity of the Modified Conconi Test for Determining Ventilatory Threshold During On-Water Rowing. J Sports Sci Med. 2011;10(4):616–623.

7. Jones AM, Doust JH. Lack of reliability in Conconi’s heart rate deflection point. Int J Sports Med. 1995;16(8):541–544. doi:10.1055/s-2007-973051

8. Hofmann P, Pokan R, Preidler K, et al. Relationship between heart rate threshold, lactate turn point and myocardial function. Int J Sports Med. 1994;15(5):232–237. doi:10.1055/s-2007-1021052

9. Hofmann P, Pokan R, Von Duvillard SP, Seibert FJ, Zweiker R, Schmid P. Heart rate performance curve during incremental cycle ergometer exercise in healthy young male subjects. Medicine & Science in Sports & Exercise. 1997;29(6):762.

10. Kjertakov M, Dalip M, Hristovski R, Epstein Y. Prediction of lactate threshold using the modified Conconi test in distance runners. Physiol Int. 2016;103(2):262–270. doi:10.1556/036.103.2016.2.12

11. Carey D. Assessment of the accuracy of the Conconi test in determining gas analysis anaerobic threshold. J Strength Cond Res. 2002;16(4):641–644.

12. Jones AM, Doust JH. The Conconi test in not valid for estimation of the lactate turnpoint in runners. J Sports Sci. 1997;15(4):385–394. doi:10.1080/026404197367173

13. Artiga Gonzalez A, Bertschinger R, Brosda F, Dahmen T, Thumm P, Saupe D. Kinetic analysis of oxygen dynamics under a variable work rate. Hum Mov Sci. 2017;In press. doi:10.1016/j.humov.2017.08.020

14. Mongin D, Uribe A, Gateau J, et al. Dynamical analysis in a self-regulated system undergoing multiple excitations: first order differential equation approach. arXiv:190104915 [stat]. December 2018. http://arxiv.org/abs/1901.04915. Accessed February 20, 2019.

15. Bodner ME, Rhodes EC. A review of the concept of the heart rate deflection point. Sports Med. 2000;30(1):31–46. doi:10.2165/00007256-200030010-00004

16. Lambert MI, Mbambo ZH, Gibson ASC. Heart rate during training and competition for longdistance running. Journal of Sports Sciences. 1998;16(sup1):85–90. doi:10.1080/026404198366713

17. Wyss CR, Brengelmann GL, Johnson JM, Rowell LB, Niederberger M. Control of skin blood flow, sweating, and heart rate: role of skin vs. core temperature. Journal of Applied Physiology. 1974;36(6):726–733. doi:10.1152/jappl.1974.36.6.726

18. Savin WM, Davidson DM, Haskell WL. Autonomic contribution to heart rate recovery from exercise in humans. J Appl Physiol Respir Environ Exerc Physiol. 1982;53(6):1572–1575. doi:10.1152/jappl.1982.53.6.1572

19. Ferguson B. ACSM’s Guidelines for Exercise Testing and Prescription 9th Ed. 2014. J Can Chiropr Assoc. 2014;58(3):328.

20. Hawley JA, Noakes TD. Peak power output predicts maximal oxygen uptake and performance time in trained cyclists. European Journal of Applied Physiology and Occupational Physiology. 1992;65(1):79–83. doi:10.1007/bf01466278

21. Wasserman K, Whipp BJ, Koyl SN, Beaver WL. Anaerobic threshold and respiratory gas exchange during exercise. J Appl Physiol. 1973;35(2):236–243. doi:10.1152/jappl.1973.35.2.236

22. Baker JS, McCormick MC, Robergs RA. Interaction among Skeletal Muscle Metabolic Energy Systems during Intense Exercise. Journal of Nutrition and Metabolism. doi:10.1155/2010/905612

23. Gastin PB. Energy system interaction and relative contribution during maximal exercise. Sports Med. 2001;31(10):725–741. doi:10.2165/00007256-200131100-00003

24. Wells GD, Selvadurai H, Tein I. Bioenergetic provision of energy for muscular activity. Paediatr Respir Rev. 2009;10(3):83–90. doi:10.1016/j.prrv.2009.04.005

25. Christensen NJ, Galbo H. Sympathetic Nervous Activity During Exercise. Annual Review of Physiology. 1983;45(1):139–153. doi:10.1146/annurev.ph.45.030183.001035

26. Kjaer M. Epinephrine and some other hormonal responses to exercise in man: with special reference to physical training. Int J Sports Med. 1989;10(1):2–15.

27. McArdle WD, Katch FI, Katch VL. Exercise Physiology: Nutrition, Energy, and Human Performance. Lippincott Williams & Wilkins; 2010.

28. Brooks GA, Mercier J. Balance of carbohydrate and lipid utilization during exercise: the “crossover” concept. J Appl Physiol. 1994;76(6):2253–2261. doi:10.1152/jappl.1994.76.6.2253

29. Stallknecht B, Vissing J, Galbo H. Lactate production and clearance in exercise. Effects of training. A mini-review. Scand J Med Sci Sports. 1998;8(3):127–131.

30. Debold EP, Beck SE, Warshaw DM. Effect of low pH on single skeletal muscle myosin mechanics and kinetics. Am J Physiol, Cell Physiol. 2008;295(1):C173–179. doi:10.1152/ajpcell.00172.2008

31. Pokan R, Hofmann P, von Duvillard SP, et al. The heart rate turn point reliability and methodological aspects. Med Sci Sports Exerc. 1999;31(6):903–907.

